# Cellular mechanisms underlying the pro-cognitive effects of serotonin 5-HT7 receptors in a mouse model of schizophrenia

**DOI:** 10.64898/2025.12.12.693910

**Authors:** Thomas Gener, Sara Hidalgo, Cristina López-Cabezón, Carmen Arroyo-Portela, M. Victoria Puig

## Abstract

Serotonin 5-HT7 receptors (5-HT7Rs) have emerged as promising targets for treating cognitive and affective disturbances in schizophrenia and other neuropsychiatric disorders, yet their cellular substrates and circuit-level mechanisms remain poorly defined. Here, we combined immunohistochemistry, multisite electrophysiology, and behavioral assays to investigate how 5-HT7Rs modulate hippocampal–prefrontal pathways in healthy mice and in a subchronic phencyclidine (sPCP) model of cognitive and negative symptoms in schizophrenia. We found that 5-HT7Rs are abundantly expressed in both excitatory and inhibitory neurons of the dorsal hippocampus (dHPC) and medial prefrontal cortex (mPFC), with high co-expression in PV⁺ and SST⁺ interneurons (∼80% in dHPC; 55% PV⁺ and 75% SST⁺ in mPFC). In healthy mice, systemic 5-HT7R activation with the agonist AS-19 suppressed neuronal activity and synchrony within dHPC (CA1) – mPFC (PL) pathways, reducing theta and high-gamma power, theta–gamma coupling in CA1, theta coherence, and CA1→PL directional connectivity, consistent with recruitment of inhibitory microcircuits. The similarity between 5-HT7R-mediated inhibition and the circuit effects we previously described for 5-HT1AR activation, together with evidence for 5-HT7R–5-HT1AR heterodimerization, suggests that these receptors act in concert to dynamically constrain hippocampal–prefrontal circuits. sPCP treatment induced persistent recognition-memory impairments, heightened anxiety-like behavior, and pathological high-frequency synchronization of hippocampal–prefrontal networks. Blockade of 5-HT7Rs with SB-269970 or the atypical antipsychotic lurasidone (but not lurasidone combined with AS-19) rescued memory performance, reduced anxiety-like behavior, and normalized aberrant high-frequency hypersynchrony, while enhancing CA1→PL theta signaling immediately before memory acquisition. Together, these findings indicate that 5-HT7R activation exerts potent inhibitory control over hippocampal–prefrontal pathways, likely via PV and SST interneurons, and suggest that 5-HT7R blockade constitutes a promising therapeutic strategy to restore excitation–inhibition balance and enhance neural communication within brain circuits crucial for cognition and mood regulation in neuropsychiatric disorders.

## INTRODUCTION

The serotonin 5-HT7 receptor (5-HT7R) is implicated in a wide range of physiological and behavioral functions, including cognitive processes, mood regulation, nociception, sleep, and circadian rhythm control (Hagan et al., 2000; Leopoldo et al., 2011; Westrich et al., 2015). They are Gs protein-coupled excitatory receptors that have garnered attention for their complex and dynamic role in regulating brain development and plasticity, opening doors to potential therapeutic applications in neuropsychiatric, neurodegenerative, and neurological disorders such as schizophrenia, depression, Alzheimer disease, chronic pain, and epilepsy (Aghajanian and Andrade, 2000).

Although the widespread distribution of 5-HT7Rs throughout the central nervous system is well documented, with high levels in the hypothalamus, thalamus and hippocampus (HPC) across species (Hedlund, 2009; Leopoldo et al., 2011; Leiser et al., 2015), their cell-type-specific expression within distinct neuronal populations in health and disease remains largely uncharacterized. This gap in knowledge is particularly relevant to brain regions critically implicated in neuropsychiatric disorders, such as the HPC and the prefrontal cortex (PFC), whose pathophysiology is closely linked to cognitive and affective dysfunction.

Preclinical studies have consistently implicated 5-HT7Rs in brain development and neural plasticity (Beique et al., 2004; Meneses, 2017; Crispino et al., 2020; Olusakin et al., 2020). Notably, the selective antagonist SB-269970 has demonstrated strong pro-cognitive effects in preclinical models of N-methyl-D-aspartate receptor (NMDAR) hypofunction, a pathophysiological mechanism relevant to schizophrenia (McLean et al., 2009; Bonaventure et al., 2011; Waters et al., 2012; Meltzer et al., 2013; Adell, 2025). These findings position 5-HT7Rs as compelling targets for therapeutic strategies aimed at cognitive impairment, particularly in disorders such as schizophrenia.

Atypical antipsychotics (APDs) such as lurasidone, risperidone, clozapine, and aripiprazole act as 5-HT7R antagonists, with varying degrees of affinity (Roth et al., 1994; Fukuyama et al., 2023). Preclinical evidence suggests that blockade of 5-HT7Rs by APDs may enhance cognitive flexibility, working memory, and mood regulation, domains often impaired in schizophrenia and poorly addressed by traditional dopaminergic approaches (Meltzer and Massey, 2011; Meneses, 2015). Among atypical APDs, lurasidone shows a remarkable affinity for 5-HT7Rs (Ki 0.5 nM), while it also acts as an antagonist at dopamine D2Rs and serotonin 5-HT2ARs and shows partial agonism at 5-HT1ARs. Its minimal affinity for receptors associated with sedation and metabolic side effects, such as H1, further distinguishes lurasidone among atypical APDs (Okubo et al., 2021; Fukuyama et al., 2023). This receptor binding profile suggests that lurasidone may be particularly suitable for addressing cognitive and mood-related deficits in schizophrenia.

Accordingly, lurasidone significantly improves negative and depressive symptoms in patients with schizophrenia, improves quality of life and social functioning relative to placebo, and offers an excellent safety and metabolic profile (Huhn et al., 2019).

In this study, we first assessed the expression of 5-HT7Rs in two key populations of inhibitory interneurons, parvalbumin (PV) and somatostatin (SST) cells, within the dHPC and mPFC, both of which are critical for network oscillations and inter-regional communication. Moreover, we performed a comprehensive analysis of how pharmacological 5-HT7R activation with the selective agonist AS-19 modulates hippocampal-prefrontal neural dynamics under physiological conditions in freely behaving mice. We then evaluated the therapeutic effects of SB-269970 and lurasidone in mitigating memory deficits and anxiety in an NMDA receptor hypofunction model of schizophrenia, using subchronic phencyclidine (sPCP) administration (Yuen et al., 2012; Horisawa et al., 2013; Meltzer et al., 2013; Rajagopal et al., 2016; Fukuyama et al., 2023), and investigated the neural substrates of the rescue (Delgado-Sallent et al., 2022, 2023; Puig et al., 2025).

Through this multi-level approach, we sought to elucidate the cellular and circuit mechanisms by which 5-HT7Rs influence cognition and negative symptoms, thereby advancing our understanding of their therapeutic potential in schizophrenia and related psychiatric disorders.

## RESULTS

### High expression of 5-HT7 receptors in parvalbumin and somatostatin interneurons within hippocampal-prefrontal circuits

Widespread 5-HT7R immunoreactivity was observed in both the dHPC and the mPFC, consistent with previous reports (Bickmeyer et al., 2002; Leiser et al., 2015). In the dHPC, 5-HT7Rs were abundantly expressed by granule cells in the CA1 and CA3 regions. In the mPFC, moderate to high 5-HT7R expression was detected in the anterior cingulate cortex (ACC), the prelimbic cortex (PL), and the infralimbic cortex (IL), with strongest labeling in cortical layers 2/3 and lower levels in layers 5 and 6 (Figure 1a).

**Figure 1.**
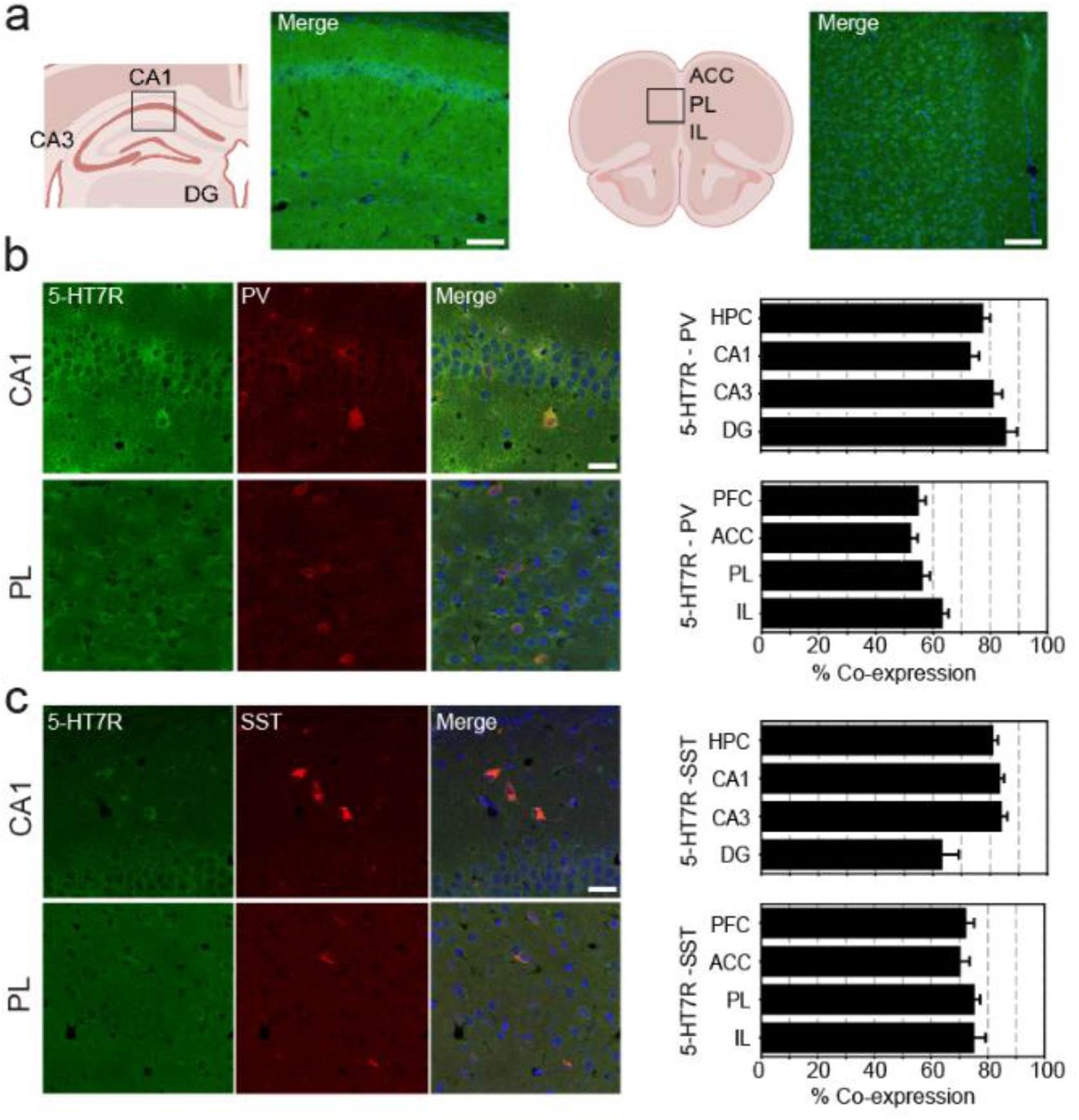
High expression of 5-HT7Rs in PV^+^ and SST^+^ interneurons of the dHPC and mPFC. **(a)** Overview of 5-HT7R expression in the CA1 dHPC and PL mPFC. Images acquired at 20x magnification; scale bar = 100 μm. **(b)** Representative confocal images showing 5-HT7R expression (green) in PV^+^ interneurons (red) with DAPI nuclear counterstaining (blue) in CA1 and PL. Quantification of 5-HT7R and PV co-expression across subregions of the dHPC and mPFC is also shown. Images acquired at 63x magnification; scale bar = 25 μm. **(c)** Same as in (b), but showing co-expression of 5-HT7Rs (green) with SST^+^ interneurons (red) instead of PV^+^. Quantification of 5-HT7R and SST co-expression is presented for all regions analyzed. See Table 1 for detailed quantification.

**Table 1.** Expression of 5-HT7Rs in PV^+^ and SST^+^ interneurons across subregions of the dHPC and mPFC. The same animals were used for both immunohistochemical assays, with interleaved brain sections analyzed for each marker. \**p* < 0.05, male vs. female (unpaired Student’s *t*-test).

| Sub-region | Sex | Number of animals | Mean PV<sup>+</sup> Count per Section | % Co-expression | Number of animals | Mean SST<sup>+</sup> Count per Section | % Co-expression |
| --- | --- | --- | --- | --- | --- | --- | --- |
| CA1 | Female | 4 | 27±4 | 72±6 | 3 | 16±4 | 83±7 |
|  | Male | 4 | 32±4 | 75±8 | 3 | 19±3 | 83±6 |
|  | All | 8 | 30±3 | 74±5 | 6 | 18±2 | 83±4 |
| CA3 | Female | 4 | 23±3 | 80±4 | 3 | 10±2 | 86±8 |
|  | Male | 4 | 27±4 | 83±8 | 3 | 13±2 | 82±6 |
|  | All | 8 | 25±2 | 82±4 | 6 | 12±2 | 84±5 |
| DG | Female | 4 | 7±1 | 83±9 | 3 | 3±1 | 63±26 |
|  | Male | 4 | 8±3 | 88±8 | 3 | 4±2 | 63±17 |
|  | All | 8 | 7±2 | 86±6 | 6 | 4±1 | 63±15 |
| ACC | Female | 4 | 101±6 | 49±3 | 3 | 44±4 | 74±5 |
|  | Male | 3 | 117±10 | 57±3 | 3 | 41±7 | 67±7 |
|  | All | 7 | 108±6 | 52±3 * | 6 | 43±4 | 71±4 |
| PL | Female | 4 | 65±7 | 52±5 | 4 | 33±3 | 77±5 |
|  | Male | 3 | 74±7 | 62±3 | 3 | 28±8 | 73±2 |
|  | All | 7 | 69±5 | 56±3 | 7 | 31±3 | 76±3 |
| IL | Female | 4 | 32±6 | 61±6 | 4 | 7±2 | 81±10 |
|  | Male | 3 | 32±7 | 66±5 | 3 | 11±3 | 72±7 |
|  | All | 7 | 32±5 | 63±4 | 7 | 8±2 | 78±7 * |

We next examined 5-HT7R expression in two key interneuron populations, parvalbumin-expressing (PV^+^) and somatostatin-expressing (SST^+^) cells, which are known to play essential roles in both local and long-range neural dynamics. In the dHPC, 5-HT7Rs were highly expressed in both PV^+^ and SST^+^ interneurons, with approximately 80% of cells showing immunoreactivity across CA1, CA3, and DG (n = 8 mice). In the mPFC, 5-HT7Rs were expressed by 55% of PV^+^ neurons and 75% of SST^+^ neurons across the ACC, PL, and IL subregions (Figure 1b,c; Table 1). Notably, 5-HT7R co-expression with PV^+^ and SST^+^ interneurons was comparable between female and male mice, with only minor regional differences. Specifically, 5-HT7R-PV co-expression was slightly higher in the ACC of males, whereas 5-HT7R-SST co-expression was elevated in the IL cortex of females (Student’s *t*-test, *p* < 0.04; Table 1).

These findings reveal robust 5-HT7R expression in PV^+^ and SST^+^ interneurons in both brain regions, with overall higher co-expression levels in the dHPC compared to the mPFC.

### Activation of 5-HT7Rs suppresses neuronal activity and synchrony within hippocampal–prefrontal circuits

We next investigated the role of 5-HT7Rs in the regulation of physiological neural dynamics within the CA1 region of the dHPC and the PL mPFC, a circuit critically implicated in cognitive function and consistently impaired in schizophrenia. To this end, we recorded neural activity in both regions in freely moving mice, before and after the administration of saline, the selective full agonist AS-19 (10 mg/kg, i.p.), and the selective antagonist SB-269970 (two injections of 4 mg/kg, i.p.) at doses previously validated in rodents (Fukuyama et al., 2023). We specifically assessed changes in key electrophysiological markers of circuit function, including oscillatory power, cross-frequency phase–amplitude coupling (PAC), multi-unit spiking activity (MUA), interregional phase coherence (wPLI), and directional connectivity (phase slope index, PSI). Our previous work has comprehensively characterized the hippocampal-prefrontal circuit, showing that it exhibits intrinsic synchronization within the theta (8-10 Hz) and gamma (30-100 Hz) frequency bands under physiological conditions (Puig et al., 2025). The present approach extends this framework by examining serotonergic modulation of hippocampal–prefrontal synchrony through 5-HT7Rs, building upon our prior investigations of 5-HT1A, 5-HT2A, and 5-HT4 receptors signaling in freely behaving mice (Gener et al., 2019, 2025; Puig et al., 2025).

Activation of 5-HT7Rs by the selective agonist AS-19 disrupted theta-band oscillations in CA1 and PL cortex (n = 7 male mice; rmANOVA; F_2,12_ = 5.275 and 3.829, p = 0.023 and 0.05, respectively), which was accompanied by spectral power increases toward the adjacent frequency bands, delta (2–5 Hz) and alpha (10–14 Hz). In addition, high gamma oscillations (52–100 Hz) were significantly reduced in both CA1 and PL (F_2,12_ = 5.044, 6.443; p = 0.026, 0.013, respectively), with the largest reductions observed in CA1 (Figure 2a,b). Consistent with these findings, AS-19 administration disrupted theta–gamma coupling in CA1 (phases between 5 and 10 Hz and amplitudes between 50 and 100 Hz; F_2,12_ = 38.414, p < 0.001; Figure 2c,d). Moreover, a small decrease in multi-unit spiking activity was detected in both regions (CA1, PL: F_2,10_ = 9.187, 14.252, p = 0.005, 0.001; Figure 2e).

**Figure 2.**
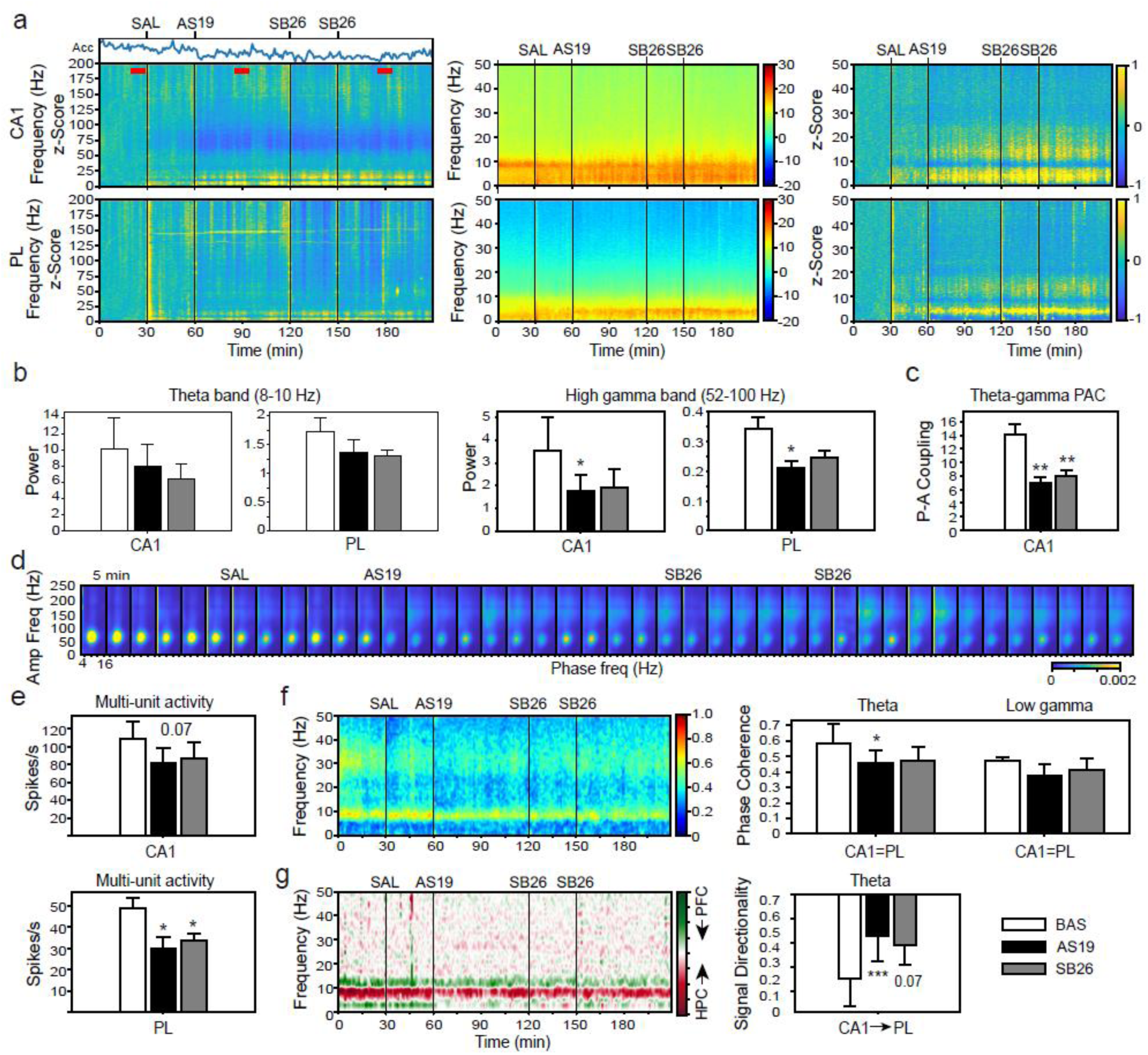
Selective 5-HT7R activation inhibits neural activity and synchrony of hippocampal – prefrontal circuits. **a**) Average power spectrograms (power per minute; n = 7 mice) and corresponding z-scores of local field potentials recorded in CA1 and PL cortex following administration of saline, the 5-HT7R agonist AS-19 (AS19; 10 mg/kg, i.p.) and the 5-HT7R antagonist SB-269970 (SB26; 4 + 4 mg/kg, i.p.). Quantification of locomotor activity is shown as the variance of the accelerometer signal per minute (Acc). Red lines indicate 10-min quantification epochs used in the panels below. **b**) Quantification of theta and high-gamma power across drug conditions. Units in mV^2^/Hz. **c,d**) Quantification and average effects of the compounds on theta-high gamma comodulation in CA1 (phase: 5-10 Hz, amplitude: 50-100 Hz). **e**) Effects of the compounds on multi-unit activity (MUA) recorded in both regions. **f**) Phase coherence between CA1 and PL and corresponding quantification of the theta (8-10 Hz) and low gamma (25-40 Hz) bands. **g**) Signal directionality between CA1 and PL and corresponding quantification at theta (6-10 Hz). \**p* < 0.05, \*\**p <* 0.01, and \*\*\**p <* 0.001 relative to baseline (BAS; rmANOVA followed by Holm’s *post-hoc* test). Data are mean ± S.E.M.

At the circuit level, CA1-PL phase coherence was significantly weakened at theta frequencies (F_2,10_ = 4.211; p = 0.047), while the low gamma band (25-40 Hz) showed a non-significant trend toward decrease (Figure 2f). Additionally, directed information flow from CA1 to PL within the theta range (6-10 Hz) was strongly attenuated (F_2,6_ = 21.759, p = 0.002; Figure 2g). Administration of SB-269970 did not reverse most AS-19-induced neural alterations (Figure 2a-g), with the exception of a partial recovery of high gamma oscillations and theta-band directional signaling from CA1 to PL (Figure 2b,g).

Moreover, AS-19 produced reductions in locomotor activity (Acc: F_2,12_ = 10.499, p = 0.002; Figure 2a), although animals remained actively exploring throughout the recording sessions, indicating that the neural effects cannot be attributed solely to reduced movement.

These results support a strong inhibitory role of 5-HT7R activation in hippocampal–prefrontal local and circuit dynamics. Because 5-HT7Rs are Gs-coupled excitatory receptors and induce excitatory responses in pyramidal neurons of the CA1 and PL cortex (Fan et al., 2011; Siwiec et al., 2020), we hypothesized that feedforward inhibition driven by 5-HT7R-expressing interneurons contributes critically to the cellular mechanisms of these receptors.

### 5-HT7R antagonism with SB-269970 and lurasidone rescues short- and long-term recognition memory deficits and reduces anxiety in a model of schizophrenia

We evaluated the memory-enhancing properties of SB-269970 and lurasidone at doses previously shown to exert pro-cognitive effects (Rajagopal et al., 2014, 2016; Huang et al., 2018). Recognition memory was assessed using the novel object recognition (NOR) test, following protocols established in our earlier studies (Alemany-González et al., 2020, 2022; Delgado-Sallent et al., 2023; Gener et al., 2025). Short-term (STM, 3 mins) and long-term (LTM, 24 h) object recognition memory were tested under baseline conditions and after subchronic phencyclidine (sPCP) administration (10 injections, 10 mg/kg, s.c.; Figure 3a,b), a well-validated model of schizophrenia-related cognitive deficits (Antunes and Biala, 2011; Rajagopal et al., 2014). Mice (n = 8 males) received saline or SB-269970 (4 mg/kg, i.p.) one hour before the familiarization phase in sessions conducted one week apart, both before and after sPCP treatment. This design aimed to potentiate memory acquisition through 5-HT7R-mediated pro-cognitive modulation. During weeks 3 and 4 post-sPCP, we additionally assessed the memory-rescuing effects of the atypical antipsychotic lurasidone (LUR, 0.6 mg/kg, i.p.) and the combination of lurasidone and the 5-HT7R agonist AS-19 (10 mg/kg, i.p.; Huang et al., 2012; Rajagopal et al., 2014). Finally, saline (n = 5) and SB-269970 (n = 2) were re-tested during weeks 6 and 7 post-sPCP (Figure 3a). Neural activity in CA1 and the PL cortex was simultaneously recorded during the one-hour period preceding the familiarization phase (Figure 3b).

**Figure 3.**
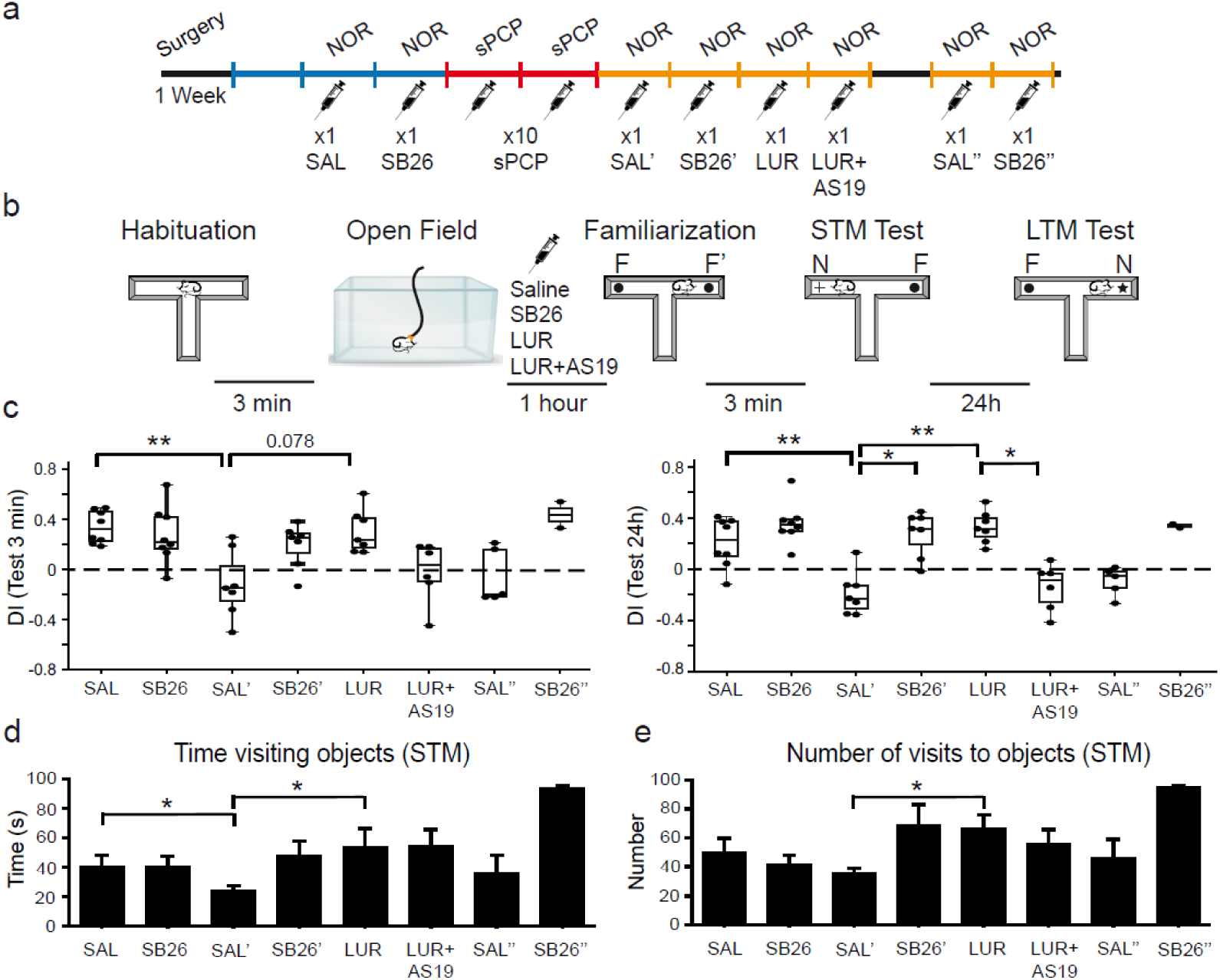
Blockade of 5-HT7Rs rescues recognition memory and reduces anxiety in the sPCP model of schizophrenia. **(a)** Experimental design. Mice were surgically implanted with recording electrodes in CA1 and the PL cortex. During the first recovery week, animals were handled and habituated to the recording setup and cable. Saline and SB-269970 (SB26; 4 mg/kg, i.p.) were administered immediately before the familiarization phase in two consecutive weeks, conducted both before and after sPCP treatment (10 mg/kg, s.c.). During weeks 3 and 4 post-sPCP, mice received either lurasidone (LUR; 0.6 mg/kg, i.p.) or a combination of lurasidone and AS-19 (LUR-AS19; 10 mg/kg). Finally, during weeks 6 and 7 post-sPCP, animals were retested with saline (n = 5 mice) and SB-269970 (n = 2) to assess persistence of cognitive and anxiety-related effects. **(b)** The NOR task consisted of four 10-min phases. Electrophysiological recordings were conducted in an open field arena the hour preceding the familiarization phase. **(c)** Drug effects on discrimination indices (DIs) of the 3-minute (STM) and 24h (LTM) memory tests. **(d,e)** Quantification of object exploration time and number of visits across treatment groups during the 10-minute testing sessions. rmANOVA followed by Holm’s *post hoc* test, \**p* < 0.05, ** *p* < 0.01. Data are mean ± S.E.M.

Mice exhibited positive discrimination indices (DIs; calculated as [time exploring the novel object - time exploring the familiar object] / total exploration time) following saline administration at both the 3-minute (STM) and 24-hour (LTM) memory intervals (0.34 ± 0.05 and 0.21 ± 0.07, respectively), confirming good recognition memory under baseline conditions. Administration of SB-269970 did not improve memory performance in either test, as DIs did not differ significantly from those obtained under saline (rmANOVA SAL vs. SB26, *p* > 0.05; Figure 3c).

As anticipated, sPCP treatment produced a marked impairment in recognition memory, evidenced by significant reductions in both STM and LTM DIs compared with baseline saline performance (paired *t*-test, [SAL, SAL’]; STM, LTM: *p* = 0.003 and 0.004, respectively). Treatment with either SB-269970 or lurasidone effectively rescued STM and LTM performance in most animals, resulting in significant group-level improvements in DI values (rmANOVA, [SAL’, SB26’, LUR]; F_2,8_ = 9.02 and 15.923, *p* = 0.009 and 0.002, respectively). The memory-enhancing effects of lurasidone were likely mediated via 5-HT7R blockade, as co-administration of lurasidone with AS-19 failed to restore STM or LTM performance ([SAL’, LUR, LUR-AS19]; F_2,8_ = 6.489 and 13.127, *p* = 0.021 and 0.003; Figure 3c). Importantly, recognition memory deficits persisted for at least six weeks after sPCP exposure (n = 5 mice), yet remained responsive to pharmacological rescue by SB-269970 (n = 2 mice; Figure 3c).

We next examined whether 5-HT7R modulation produced anxiolytic effects in the sPCP model during NOR testing, in line with our previous observations of anxiolysis induced by 5-HT4R agonism (Gener et al., 2025). Following sPCP treatment, mice exhibited heightened anxiety-like behavior during the STM task, evident in their increased reactivity to drug injection and experimenter handling prior to testing. This was reflected by a marked reduction in object exploration, as animals spent significantly less time visiting the two objects compared with their pre-sPCP baseline (paired *t*-test, [SAL vs. SAL’]; 23 ± 4 s vs. 42 ± 8 s; *p* = 0.018; *n* = 7). In fact, after sPCP exposure, object exploration following saline injection was minimal, likely reflecting avoidance of the brightly illuminated maze containing a novel object. In contrast, administration of SB-269970 or lurasidone significantly increased exploration time under the same conditions (rmANOVA, [SAL’, SB26’, LUR]; F_2,8_ = 6.845; *p* = 0.019; *n* = 5), which was accompanied by a higher number of object visits (F_2,8_ = 7.362, *p* = 0.015). These anxiolytic-like effects were replicated during weeks 6 and 7 post-sPCP, indicating a persistent responsiveness to 5-HT7R antagonism despite the long-lasting behavioral alterations induced by sPCP (Figure 3d,e).

Collectively, these findings indicate that both SB-269970 and lurasidone attenuate the mnemonic and anxiety-like deficits induced by sPCP administration.

### 5-HT7R antagonism mitigates sPCP-induced hypersynchronization and enhances connectivity in hippocampal-prefrontal circuits

In the preceding behavioral experiments, we recorded neural activities simultaneously from the CA1 and PL subdivisions during the hour preceding object familiarization. Following a 15-minute baseline, animals received saline (n = 5), SB-269970 (4 mg/kg, i.p.; n = 7), lurasidone (0.6 mg/kg, i.p.; n = 6), or a combination of lurasidone and AS-19 (0.6 mg/kg and 10 mg/kg, i.p.; n = 5). Recordings continued for one hour post-injection until the onset of the familiarization phase (Figure 3b).

Under healthy conditions, administration of saline and SB-269970 did not alter local or long-range synchrony within hippocampal-prefrontal circuits. In contrast, following sPCP exposure, saline injections uncovered a pathological hypersynchronization in both CA1 and PL regions, characterized by abnormally high-frequency oscillations (150-200 Hz) that were not present under baseline conditions (rmANOVA [SAL, SAL’]; CA1, PL: F_3,19_ = 6.251 and 6.28, *p* = 0.034 and 0.035, respectively). This aberrant activity was most prominent in the PL cortex, extending above 100 Hz and beyond 200 Hz, while in CA1, it emerged primarily above 150 Hz (Figure 4a). These abnormal high-frequency rhythms were strongly suppressed by SB-269970 and lurasidone, but further exacerbated by co-administration of lurasidone and AS-19 (Figure 4c,e; [SAL’, SB26’, LUR, LUR+AS-19]; CA1, PL: F_3,19_ = 7.3 and 5.89, *p* = 0.014 and 0.025, respectively).

**Figure 4.**
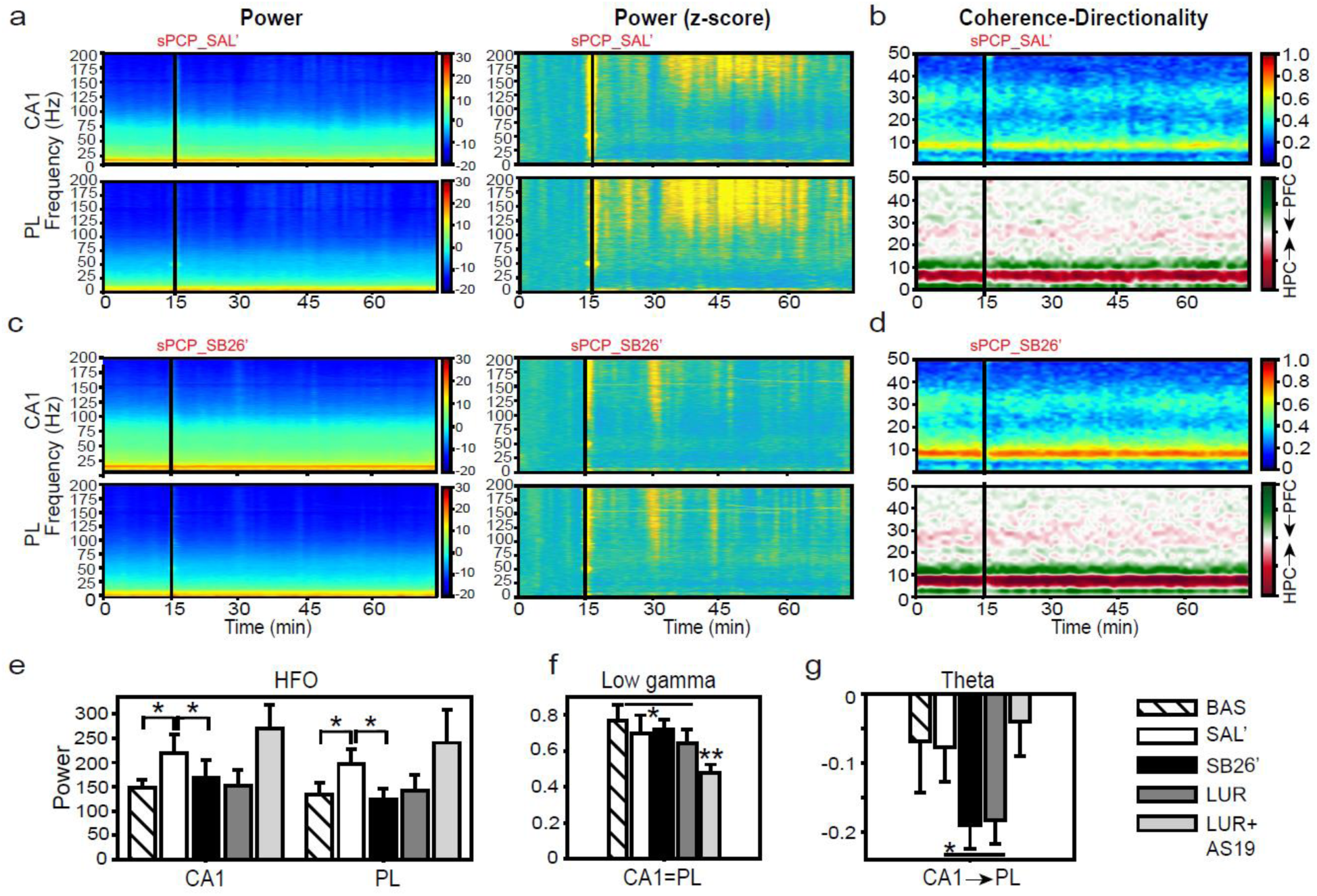
5-HT7R antagonism reduces sPCP-induced hypersynchronization and enhances hippocampal-prefrontal connectivity. **(a)** Average power spectrograms (power per minute; n = 5 mice) and corresponding z-scores of local field potentials recorded in CA1 and PL cortex during the hour preceding the NOR familiarization phase following saline administration (sPCP-SAL’). Both regions exhibited pathological hypersynchronization at high frequencies (>100 Hz). **(b)** A disruption in the low gamma coherence band (25-40 Hz) was also observed (top) while no overt changes in signal directionality were appreciated (bottom). **(c)** Treatment with SB-269970 effectively suppressed high-frequency activity in the CA1 and PL regions (n = 7 mice). **(d)** Coherence was not altered by SB-269970 (top) but theta CA1-to-PL directionality (6-10 Hz) was strengthened. **(e)** Quantification of high-frequency power (150-200 Hz) across experimental groups. Units in mV^2^/Hz. **(f)** Quantification of low gamma (25-40 Hz) CA1-PL coherence across experimental groups. **(g)** Directed theta band (6-10 Hz) information flow from CA1 to PL was weakened after saline, enhanced by SB-269970 and lurasidone, and further decreased by lurasidone + AS-19. rmANOVA followed by Holm’s *post hoc* test, \**p* < 0.05, ** *p* < 0.01. Data are mean ± S.E.M.

Notably, when retested at week 6, saline again elicited elevated high-frequency power, which was significantly reduced by SB-269970 a week later (CA1, PL: F_1,7_ = 8.529 and 5.346, *p* = 0.022 and 0.05, respectively). Theta-gamma phase-amplitude coupling in CA1 remained unchanged across treatments.

At the circuit level, CA1-PL coherence in the low gamma band (25-40 Hz) was significantly reduced following saline administration, and this disruption was not reversed by either SB-269970 or lurasidone, but was further exacerbated by co-administration of lurasidone with AS-19 (F_3,19_ = 7.172, *p* = 0.015; Figure 4b,d,f). Conversely, directed theta-band (6-10 Hz) information flow from CA1 to PL was enhanced by SB-269970 lurasidone relative to saline, and further weakened by lurasidone + AS-19 (F_3,17_ = 4.35, p = 0.05; Figure 4b,d,g).

Collectively, these findings demonstrate that 5-HT7R antagonism counteracts sPCP-induced network hypersynchronization and enhances hippocampal-to-prefrontal information flow, thereby restoring excitatory-inhibitory balance and functional connectivity within the circuit immediately prior to memory acquisition.

## DISCUSSION

In this study, we investigated how serotonin 5-HT7Rs shape hippocampal-prefrontal pathways under physiological conditions and following NMDAR hypofunction in the sPCP model of schizophrenia. Three main conclusions emerge from our findings. First, 5-HT7Rs are abundantly expressed in both excitatory and inhibitory PV⁺ and SST⁺ interneurons within the dHPC and mPFC, positioning this receptor as a major regulator of inhibitory microcircuits. Second, systemic activation of 5-HT7Rs with the selective agonist AS-19 exerts a potent inhibitory influence on dHPC-mPFC neural activity and synchrony, suppressing local oscillatory power, phase-amplitude coupling, long-range coherence, and directional connectivity, consistent with potent recruitment of feedforward inhibition. Third, pharmacological blockade of 5-HT7Rs with SB-269970 and lurasidone, but not lurasidone combined with AS-19, restored recognition memory, reduced anxiety-like behavior, and normalized pathological high-frequency hypersynchronization, while enhancing CA1→PL theta signaling immediately prior to memory acquisition, linking 5-HT7R signaling to both cognitive and circuit rescue.

Our anatomical data extend previous work showing dense 5-HT7R expression in hippocampal and prefrontal networks by identifying PV⁺ and SST⁺ interneurons as relevant cellular substrates in these regions. The high prevalence of 5-HT7Rs in PV⁺ and SST⁺ cells (∼80% in dHPC; 55–75% in mPFC) suggests that 5-HT7R signaling can exert broad control over perisomatic and dendritic inhibition, and thereby over the timing, gain, and integration properties of principal neurons. The overall higher co-expression in dHPC compared with mPFC, points to region-specific modes of 5-HT7R control over inhibitory circuits. This refined cellular map provides a framework for interpreting the profound effects of 5-HT7R modulation on hippocampal–prefrontal dynamics. This distribution pattern aligns with earlier anatomical studies reporting dense 5-HT7R mRNA and protein expression in all prefrontal and hippocampal subfields (Leopoldo et al., 2011; Leiser et al., 2015), and refines recent studies reporting expression of 5-HT7Rs by PV⁺ and SST⁺ interneurons in cortex, hippocampus, and other brain regions (Pehrson et al., 2016; Kusek et al., 2021; Siwiec et al., 2024; Schmitz et al., 2025).

Functionally, 5-HT7R activation by AS-19 produced a coordinated suppression of hippocampal– prefrontal activity. We observed reductions in theta and high-gamma power, decreased CA1 theta–gamma coupling, diminished CA1–PL theta coherence, and weakened CA1→PL theta-directed connectivity, accompanied by a modest reduction in multi-unit firing. These effects are striking given that 5-HT7Rs are Gs-coupled and enhance the excitability of pyramidal neurons in CA1 and mPFC (Tokarski et al., 2003; Fan et al., 2011; Siwiec et al., 2020). Our data therefore support a model in which activation of 5-HT7Rs on PV⁺ and SST⁺ interneurons produces powerful feedforward inhibition that overrides direct excitatory actions on pyramidal cells. This interpretation is consistent with recent findings in the basal amygdala, where 5-HT7R activation depolarizes PV⁺ interneurons and strengthens inhibitory drive onto projection neurons (Kusek et al., 2021). The similarity between 5-HT7R-mediated inhibition and the circuit effects we previously described for 5-HT1AR activation (Gener et al., 2018, 2019), together with evidence for 5-HT7R–5-HT1AR heterodimerization (Naumenko et al., 2014), suggests that these receptors act in concert to dynamically constrain hippocampal–prefrontal communication.

In the sPCP model, we found that subchronic NMDAR hypofunction induced lasting impairments in short-and long-term recognition memory and heightened anxiety-like behavior during the NOR task. These effects are consistent with the cognitive and affective deficits characteristic of schizophrenia. Treatment with either SB-269970 or lurasidone restored recognition memory performance and increased object exploration in sPCP-treated mice, whereas SB-269970 had no effect in healthy animals. These findings align with previous reports highlighting the procognitive abilities of SB-269970 and lurasidone in preclinical models of NMDAR hypofunction (Bonaventure et al., 2011; Meltzer et al., 2011; Horiguchi et al., 2012; Waters et al., 2012; Horisawa et al., 2013; Rajagopal et al., 2016). Similarly, we recently reported comparable procognitive and anxiolytic effects of the 5-HT4R agonist RS-67333 in the same model (Gener et al., 2025). Consistent with these results, 5-HT7Rs in CA1 and PL cortex have been implicated in the regulation of anxiety- and depression-like behaviors in rodents (Leopoldo et al., 2011; Zhang et al., 2015; Du et al., 2018; Fukuyama et al., 2023). Critically, the fact that co-administration of lurasidone with the 5-HT7R agonist AS-19 prevented the procognitive effects strongly implicates 5-HT7R blockade as a key contributor to lurasidone’s efficacy in this paradigm.

At the circuit level, saline injections unveiled robust high-frequency hypersynchronization in both CA1 and the PL cortex of sPCP-treated mice, and were particularly pronounced in the PL region. SB-269970 and lurasidone effectively suppressed pathological high-frequency oscillations and enhanced CA1→PL theta signaling, further linking 5-HT7R antagonism to the stabilization of circuit function. These findings support a model in which 5-HT7R blockade rebalances excitation–inhibition within hippocampal–prefrontal loops, dampening aberrant high-frequency activity while preserving or restoring task-relevant synchrony. In particular, they corroborate our previous findings demonstrating a critical role of CA1→PL theta connectivity in NOR encoding (Alemany-González et al., 2020, 2022; Delgado-Sallent et al., 2023).

Several limitations should be acknowledged. First, our pharmacological approach does not allow us to disentangle the relative contributions of 5-HT7Rs expressed in PV⁺ versus SST⁺ interneurons or in pyramidal neurons and other cell types to behavior and neural dynamics. Cell type–specific manipulations (e.g., conditional knockouts, DREADDs, or optogenetic tagging of 5-HT7R-expressing interneurons) will be required to resolve these contributions. Second, we focused on male mice in the electrophysiological experiments, while the anatomical data revealed sex-dependent differences in mPFC expression (although small). Future work should systematically examine whether 5-HT7R-driven modulation of hippocampal–prefrontal dynamics and cognition is sexually dimorphic. Third, although the sPCP model captures key aspects of NMDAR hypofunction, extrapolation to the full clinical heterogeneity of schizophrenia must be made with caution, especially results related to anxiety, which were not directly investigated in this study.

In summary, our findings identify 5-HT7Rs as pivotal modulators of hippocampal–prefrontal circuit function, exerting inhibitory control under physiological conditions while becoming therapeutically relevant targets under NMDA receptor hypofunction. The dual ability of 5-HT7Rs to regulate both local synchronization and long-range inter-areal communication positions them as key nodes for maintaining excitatory–inhibitory balance in cortical–limbic networks. These results highlight 5-HT7R antagonism as a mechanistically grounded approach to alleviate cognitive and affective disturbances in schizophrenia and related brain disorders.

## Author Contributions

Conceptualization, M.V.P.; Methodology, M.V.P., T.G., S.H-N.,, C.A-P, C.L-C.; Formal Analysis, M.V.P., T.G., S.H-N., C.A-P; Investigation, M.V.P., T.G., S.H-N., C.L-C., C.A-P; Data Curation, M.V.P., T.G., S.H-N., C.A-P; Writing – Original Draft Preparation, M.V.P.; Writing – Review & Editing, M.V.P., T.G., S.H-N., C.A-P; Visualization, M.V.P., T.G., S.H-N., C.A-P; Supervision, M.V.P. and T.G.; Project Administration, M.V.P.; Funding Acquisition, M.V.P.

## MATERIALS AND METHODS

### Animals and housing

Adult female and male C57Bl6/J mice (3–4 months old, 20–30 g) were obtained from Charles River. Mice were housed under controlled environmental conditions, including a temperature of 22 ± 2 °C, relative humidity of 60%, and a 12:12 light–dark cycle (lights off from 8:00 p.m. to 20:00 a.m.). Food (standard pellet diet) and water were available *ad libitum* throughout the study, except during the brief behavioral and recording sessions, when the animals did not have access to food or water. All procedures were conducted in compliance with EU directive 2010/63/EU and Spanish guidelines (Laws 32/2007, 6/2013 and Real Decreto 53/2013) and were authorized by the University of Barcelona Animal Research Ethics Committee (project 150_24).

### Immunohistochemistry

Mice were perfused with 4% PFA and the brains post-fixed in 4% PFA for 24 hours at 4°C. Then, brains were cryoprotected by immersing them in 30% sucrose in PBS overnight at 4°C until they sank. The brains were then frozen at -80° until they were processed. Brains were later sectioned at a thickness of 30 μm using a cryostat. Brain sections containing the HPC and the mPFC (at least three sections per area) were washed in PBS at RT (3 times for 10 minutes) and later incubated with blocking solution (5% Donkey serum, 0.3% Triton X-100 and 100mM of glycine in PBS) (Rosas-Arellano et al., 2016) overnight at RT. After blocking, the tissue sections were incubated in a solution containing the primary antibodies in blocking solution over-weekend at 4°C and an extra hour at RT. The primary antibodies were: rabbit anti-5-HT7R (1:200, Invitrogen MA5-37958), sheep anti-PV (1:500, Invitrogen PA5-47693), and mouse anti-SST (1:500, GeneTex GTX71935). In the case of 5-HT7R and SST assays, the sections were incubated 60 minutes at RT with a solution of ReadyProbes™ Mouse on Mouse IgG Blocking Solution (30X) (Invitrogen R37621) and PBS and were washed in PBS at RT (3 times for 10 minutes) previous to the primary antibodies incubation to avoid nonspecific binding of antibodies to the tissue. Later, tissue sections were washed in PBS at RT (3 times for 10 minutes) and incubated with the secondary antibodies for 2 hours at RT in the dark. The secondary antibodies were: Goat anti-rabbit IgG Alexa Fluor® 488 (1:1000, Abcam AB-150077), Donkey anti sheep Alexa Fluor™ 594 (1:1000, Invitrogen A-11016) and Goat anti mouse Alexa Fluor™ 594 (1:500, Invitrogen A-11005). Finally, the slices were cleaned in PBS (3 times for 10 minutes) and mounted with Fluoromount-G™ with DAPI (Invitrogen 00-4959-52) for nuclear staining, and covered with a coverslip. The sections were examined with a 10x and 20x objective under a Dragonfly 200 confocal microscope system (Oxford Instruments Andor) using the Fusion software (version 2.4.0.14; Oxford Instruments) and with a 63x objective with a Stellaris 8 DRIVE confocal microscope system (Leica Microsystems) using the Leica Application Suite X software (version 4.6.1.27508; Leica Microsystems). Negative control experiments with no primary antibodies were conducted to confirm specificity of the stainings. The images were visualized and analyzed using ImageJ software.

### Pharmacology

The selective 5HT7R agonist AS19 and antagonist SB269970 were obtained from Merck. The antipsychotic drug lurasidone was obtained from Sigma-Aldrich (UK). Phencyclidine (non-competitive NMDAR antagonist) was obtained from Sigma-Aldrich (UK). Drugs were diluted in saline or 1% DMSO (vehicle for lurasidone) in saline when necessary and the pH corrected to be between 6-8. Drugs were first diluted in a concentrated solution and frozen at -20C. The day of the experiment, the solution was thawed and diluted before administration. Each mouse was used to test multiple drugs and at least one week of washout was left between experiments.

### Novel object recognition test (NOR)

Recognition memory was assessed using a custom-designed T-maze, as previously reported (8 cm wide × 30 cm long × 20 cm high; Alemany-González et al., 2020; Delgado-Sallent et al., 2023; Puig et al., 2025). Novel-familiar object pairs were validated as described in (Gulinello et al., 2018), and the arm in which the novel object was placed was randomized across trials. The task consisted of a habituation phase, familiarization phase, short-term memory test (STM), and long-term memory test (LTM), each lasting ten minutes. During habituation, mice explored the maze without objects. After a one-hour interval, when electrophysiological recordings were conducted in an open field, they underwent the familiarization phase, where two identical objects were placed at the end of the lateral arms. STM and LTM tests were conducted 3 minutes and 24 hours later, respectively, with one familiar object and one novel object placed in the maze. Exploratory events were timestamped using TTL pulses sent to the acquisition system, allowing precise quantification of visit number and duration. Typically, initial visits and interactions with novel objects were longer compared to later visits and interactions with familiar objects (Delgado-Sallent et al., 2023). Only tests when the mice visited both objects for at least 5 seconds were considered valid.

### Surgical procedures for electrode placement

Mice were anesthetized with 4% isoflurane and placed in a stereotaxic apparatus. Anesthesia was maintained between 0.5–2% throughout the procedure. Small craniotomies were made above the HPC and mPFC in the same hemisphere. Several micro-screws were inserted into the skull to stabilize the implant, with one positioned over the cerebellum serving as the general ground. Stereotrodes, composed of two twisted strands of 25 μm tungsten wire (Advent, UK) insulated with heat shrink tubing, were implanted unilaterally in the CA1 area of the dHPC (AP: -1.8 mm; ML: -1.3 mm; DV: -1.15 mm) and prelimbic (PL) region of the mPFC (AP: 1.5, 2.1 mm; ML: ± 0.6, 0.25 mm; DV: -1.7 mm from bregma). Neural activity was monitored during electrode placement to confirm accurate targeting of the CA1 region. Additionally, three reference electrodes were implanted in the corpus callosum and lateral ventricles (AP: 1, 0.2, -1; ML: 1, 0.8, 1.7; DV: -1.25, -1.4, -1.5, respectively). At implantation, electrode impedances ranged from 100–400 kΩ. Electrodes were secured with dental cement and connected to an adaptor for integration with the recording system. Postoperatively, mice were given at least one week to recover, during which they were closely monitored and received analgesia and anti-inflammatory treatments. Before experiments commenced, animals were handled and habituated to the recording cable. Following the completion of experiments, electrode placements were histologically verified using Cresyl violet staining, and data from misplaced electrodes were excluded from the analyses.

### Neurophysiological recordings and analyses

Electrophysiological recordings were conducted between 9:00 a.m. and 5:00 p.m., during the light phase of the housing cycle, in freely-moving mice exploring a 17 x 28 x 11 cm cage positioned within a custom-built Faraday enclosure. Animals did not have access to food or water during the recordings but had food and water available just before and after each experiment. All the recordings were carried out with the Open Ephys system at 0.1-6000 Hz and a sampling rate of 30 kHz with Intan RHD2132 amplifiers equipped with an accelerometer. The home cage was moved from the housing room to the experimental room within the animal facility where behavioral tests and electrophysiological recordings were implemented. One or two animals were recorded simultaneously in separate cages and electrophysiological setups in the same room.

Recorded signals from each electrode were filtered offline to extract multi-unit activity (MUA) and local field potentials (LFPs). MUA was estimated by first subtracting the raw signal from each electrode with the signal from a nearby referencing electrode to remove artifacts related to animal movement. Then, continuous signals were filtered between 450-6000 Hz with Python and thresholded at -3 sigma standard deviations with Offline Sorter v3 (Plexon Inc.). To obtain LFPs, signals from each electrode were detrended, notch-filtered to remove power line artifacts (50, 100, 150 and 200 Hz), decimated, and downsampled to 1kHz. Noisy electrodes detected by visual inspection from individual channel spectrograms were not used. Power spectral density results were calculated using the multi-taper method from the spectral_connectivity package in Python (time-half-bandwidth product = 5, 9 tapers, 60s sliding time window without overlap). Spectrograms were constructed using consecutive Fourier transforms (scipy.signal.spectrogram function, 60s time window, no overlap, no detrend). A 1/f normalization was applied to power spectral density results, and power spectrograms were scaled to decibels for visualization purposes. The frequency bands considered for the band-specific analyses included: delta (2-5 Hz), theta (8-10 Hz), wide theta band in PSI (6-10 Hz), beta (18-25 Hz), low gamma (30-48 Hz), low gamma band in PLI (25-40 Hz), high gamma (52-100 Hz), and high frequencies (100-200 Hz, 100-150 Hz, and 150-200 Hz).

Phase-amplitude coupling (PAC) was measured with a Python implementation of the method described in Tort et al. (phase frequencies = [0, 15] with 1 Hz step and 4 Hz bandwidth, amplitude frequencies = [10, 250] with 5 Hz step and 10 Hz bandwidth) (Tort et al., 2008). The length of the sliding window was 300s for the overview plots and 60s for the quantifications, without overlap. PAC quantification results were obtained by averaging the values of selected areas of interest in the comodulograms. Here, we focused on theta-high gamma comodulation in CA1 (phase: 5-10 Hz, amplitude: 50-100 Hz).

Prefrontal-hippocampal phase coherence was estimated via the weighted phase-lag index (wPLI, Butterworth filter of order 3), a measure of phase synchronization between areas aimed at removing the contribution of common source zero-lag effects that allowed us to estimate the synchronization between the PFC and the HPC mitigating source signals affecting multiple regions simultaneously (Stam et al., 2007; Vinck et al., 2011; Hardmeier et al., 2014; Gener et al., 2019). PLI spectra were built applying the previous function multiple times with a 1 Hz sliding frequency window (using Butterworth bandpass filters of order 3), and PLI spectrograms were generated applying the PLI spectra function over a 60s sliding window (without overlap). LFP power, PAC and PFC-HPC wPLI are also provided as z-scores with respect to baseline statistics (i.e., data is demeaned by the baseline mean and then normalized by the baseline standard deviation). In addition, we calculated the flow of information between areas with the phase slope index (PSI) with a Python translation of MATLAB’s data2psi.m (epleng = 60s, segleng = 1s) from (Nolte et al., 2008). PSI spectrograms and spectra were constructed with the same strategy as PLI plots but using a 1 or 5 Hz sliding frequency window.

During the electrophysiological recordings, we used the accelerometer’s signals to evaluate the effects of the drugs on general mobility of mice. We found that the variance of the acceleration module (Acc), which quantifies the variation of movement across the three spatial dimensions, was largest during exploration and decreased as the animals were in quiet alertness. More specifically, we calculated the instantaneous module of raw x, y and z signals from which we measured the variance of 1-minute bins (Alemany-González et al., 2020). The Acc results were presented as a ratio to the highest value in the baseline condition of the respective experiment.

All electrophysiological biomarkers were averaged across the two electrodes within each brain region.

### Statistical analyses

All statistical analyses were conducted using JASP open-source software (version 0.19.3). Differences in 5-HT7R-PV and 5-HT7R-SST co-expression between female and male mice were assessed using unpaired Student’s *t*-tests. For the analyses of electrophysiological biomarkers in Figure 2 (power, phase-amplitude coupling [PAC], multi-unit activity (MUA), weighted phase-lag index [wPLI], phase slope index [PSI]) and accelerometer measures (Acc), we used repeated-measures ANOVAs (rmANOVA) with treatment condition (baseline, AS-19, and SB-269970 second injection) as the within-subject factor. The 10-minute quantification epochs used for these analyses are detailed in Figure 2a. The behavioral and electrophysiological effects of SB-269970, lurasidone, and the lurasidone + AS-19 combination were likewise analyzed using rmANOVAs, with specific pharmacological groupings (SAL, SB26, SAL’, SB26’, LUR, and LUR+AS-19) described in the main text. When appropriate, Holm’s *post hoc* correction was applied to adjust for multiple comparisons. All analyses were performed on raw (non-normalized) data, with statistical significance set at *p* ≤ 0.05. Results reported as mean ± S.E.M.

